# SRCompiler: Translating Stella dynamic models to open-source R scripts

**DOI:** 10.1101/2024.01.08.574614

**Authors:** Enrique Morales, Cesar Bordehore, Héctor M. Pastor, David García-García

## Abstract

Stella Architect (iSee Systems) offers an object-oriented environment that facilitates the development of dynamic models. Stella, however, does not support some of the advanced mathematical tools that can be found in mathematical software such as R. Moreover, R is open source, contains numerous libraries and resources, and is used by a large community. However, building dynamic systems, which typically use a large number of differential or difference equations, are more difficult to build in R. To take advantage of both Stella and R, we have developed the SRCompiler (Stella-R Compiler), which is a translator that exports models built in Stella into R scripts. In 2012, a translator from Stella to R was developed based on plain text treatment for Stella, but it had some limitations such as scalability and error handling. Nevertheless, our approach is more versatile, scalable and expandable since it is based on compiler design, defines a formal language, and uses the latest Stella version (v. Architect).In this paper, a classical predator-prey model has been modelled in Stella to show an applied example. Following the translation of the Stella model into R, all the parameters of the model (6 in total) have been optimised to fit model output to the Hudson Bay Company’s hare-lynx time series hunting data (1847-1935).

**Software availability:**

- Name of software: SRCompiler
- SRCompiler Developers: Pastor Héctor.M, Morales Enrique
- Web page: https://github.com/hmpp91/SRCompiler
- Available: 2021
- Program language: C++
- Program size: 197 KB
- Availability and cost: Open source

## 1. Introduction

A dynamical system is a system that relates several functions that evolve in time. Usually, dynamical systems are mathematical models expressed in terms of differential equations if time is a continuous variable, or difference equations if time is a discrete variable. They are applied in a variety of disciplines ranging from biology, conservation ecology and population dynamics (Hannon and Ruth, 2014; Haefner, 2005), astronomy (Häfner, 2000), epidemiology (Bordehore et al., 2022), economics (Sterman, 2000) and environmental management (Amadei, 2021). Modelling the real world or theoretical situations requires a deep knowledge of the processes and physical and biological laws involved in the system. However, the effort of creating these real-world models is valuable to study the time evolution of the system, which provides relevant information to understand and forecast the future states of the system, and compare scenarios. A typical example of a real-world model is the weather forecasting (Richardson, 1922), and a more exotic dynamical system could be the one predicting the collapse of global society in the mid 21st century if it kept pursuing economic growth (Herrington, 2020).

Systems dynamics (not to be confused with dynamical systems) is a methodology created during the mid-1950s by Professor Jay W. Forrester of the Massachusetts Institute of Technology (Forrester, 1990). It consists of a visual representation of dynamic systems using a causal loop diagram, where the interaction of different elements of the system can be seen. In complex, multivariate systems, the implementation of systems dynamic becomes crucial. Based on this methodology, in 1985, Barry Richmond created the Stella software modelling environment for the simulation of those models (Richmond, 1985). Stella provides a visual language that allows modelling dynamical systems using boxes, arrows and similar elements. Users may not be aware of the mathematical expressions, but such graphical elements underlie differential equations of the dynamical systems. This software approaches modelling dynamical systems to users untrained in differential equations expressed with mathematical language.

In general, models depend on some parameters, which can be constant or variable in time. Such parameters describe specific characteristics of the system and must be tuned to minimise the differences between the observed data in the real world and the modelisation results. However, parameter optimization can be a complicated task that needs specific mathematical analysis tools that are not always available in Stella. A solution to overcome this impediment is to translate the Stella model to R software, where the latter has additional mathematical and statistical analytic tools that are consistently maintained and freely distributed by the active community. In this regard, Naimi and Voinov (2012) developed a translator to convert text files generated by previous Stella versions to R, named StellaR, which was based on text processing procedure to identify different components in the Stella model. This approach has some limitations such as the difficulty of expanding the capabilities of the translator, most of all, if Stella Architect software gets an update from iSee Systems, inc. To overcome these limitations, we have developed another translator from Stella to R, named SRCompiler, from a different perspective and for the latest version of Stella.

SRCompiler is based on the theory of languages and compilers, a software specialised in analysing and translating a program’s language into another. In any given language there is a set of rules that builds a proper structure to construct phrases with different meanings, and the phrases are formed with permutations of information bricks called words. Translating a language like English is quite complicated due to the huge number of possible permutations. On the other hand, a programming language following a formalism with much fewer words is easier to translate. In a compilator, the analysis of a language can be structured in different stages (Garrido et al. 2002; Aho et al. 2007). The first stage is the lexical analyser, known as *Tokenizer*. Here, the input program is divided into individual *tokens*, which is the smallest element of a program that is meaningful to the compiler. *Tokens* are equivalent to words, punctuation marks, and symbols in our language. Once the *tokens* are identified, they are passed, one at a time, to the second stage, which is the syntax analyser. This stage is known as the *Parser* and has an understanding of the language’s grammar. The *Parser* is responsible for identifying syntax errors and for translating a program without errors into internal data structures that can be interpreted or written out in another language. The lexical and the syntax stages are referred to as the compiler’s *frontend*, while the rest of the stages are referred to as the compiler’s *backend*. The advantage of this structure is that just replacing the compiler’s *backend* we could build another translator to a different high level language (different than R) or replace the entire process by an interpreter. This characteristic makes the compiler an optimal tool for translating languages, which is more flexible and adaptable than translators based on text processing procedures as StellaR (Naimi and Voinov, 2012). For example, translating a model from Stella to R needs to model and analyse the intrinsic rules of the Stella language itself. However, the text processing procedures do not have that kind of ability. Besides, compiler-based translators are more easily expandable.

In the following section, we introduce the Stella software through a simple model. Section 3 describes the SRCompiler and in Section 4, we show the application of the SRCompiler to a classic prey-predator problem based on the Hudson Bay Company’s hare-lynx time series hunting data (1847-1935). To do so we modelled the system in Stella and translated it into R using the SRCompiler. Then, we calculated the optimised parameters to fit the historical data. The last section is devoted to discussing the results.

## 2 Stella model

Models in Stella can be created in an object-oriented environment. For example, a simple equation modelling the population of a region with a limited carrying capacity *K*, birth rate *b*, and death rate *d* could be

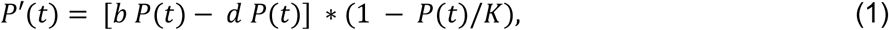

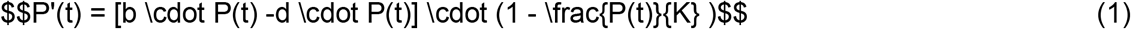

where *P(t)* is the population at time *t*, and *P’(t)* is the derivative of *P(t)*. The graphical representation of this model in Stella can be seen in Figure 1. The population is a quantifiable variable that is represented in Stella with a rectangular box called *Reservoir* or *Stock*. The variation of the population is represented by its derivative. Assuming that *b* and *d* are positive, then *P’(t)* increases as *bP(t)* increases, and decreases as *dP(t)* decreases, with both terms weighted by the term 1*-P*(*t*)*/K*, whose function is to limit the growth of the population to the carrying capacity *K*. Such limitation is always present in biological problems since the natural resources or space to sustain a population are never infinite. In Stella, increases of a reservoir are represented as *inflows* (arrows with a valve pointing to the reservoir), while decreases are *outflows* (the same valve-arrows but pointing away from the reservoir). If we represent a density-dependent system, such as a single population births and deaths model, increments of population depend on the value of the birth rate *b*, which is represented with a circumference (named *converter* or *auxiliary variable*), and a red arrow (named *connector*) pointing to the inflow. On the other hand, the decrease of population depends on the death rate *d*, which is represented in a similar way. The relationship among the reservoir, the inflows and the outflows is that a reservoir passively accumulates its inflows, minus its outflows. The clouds are stocks beyond the boundaries of the model and therefore they are not computed. The two connectors from the reservoir to the inflow and outflow valves represent the flows which fill and drain accumulations. The unfilled arrow head on the flow pipe indicates the direction of positive flow.

**Figure 1.**
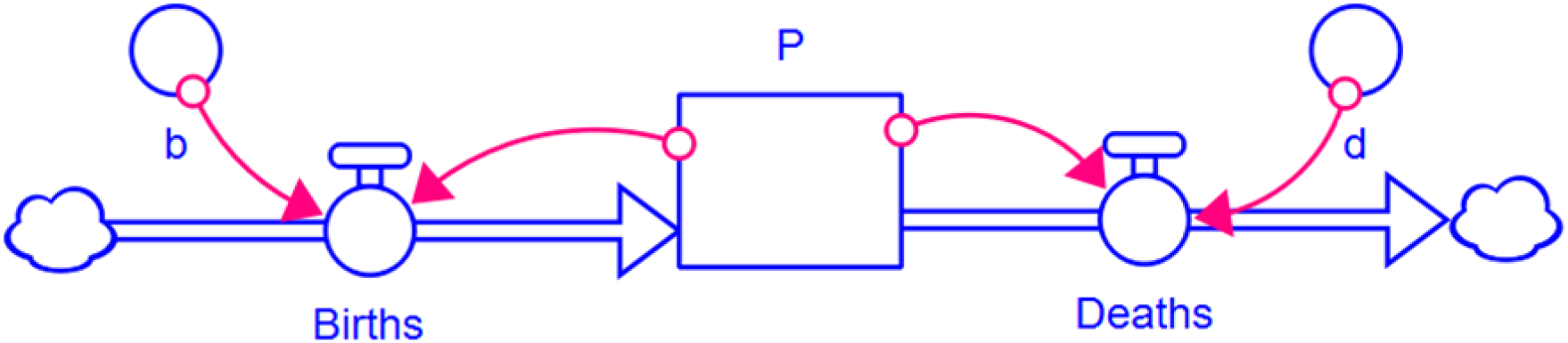
Representation in Stella of a density-dependent single population model described in Equation 1.

In a typical situation, the user introduces the visual representation shown in Figure 1 into Stella, which represents it internally as the differential equation shown in Figure 2. Such a differential equation is the mathematical model itself, which can be solved numerically by Stella by different numerical integration approaches such as Euler or Runge-Kutta 4. For example, the application of Euler with a time step of 1 unit (*dt*=1) to solve this model with coefficients *K*=12, *b*=0.02, and *d*=0.01, and the initial condition *P*(0)=0.5, is shown in Figure 3. Note that *P*(t) is always below the carrying capacity.

**Figure 2.**
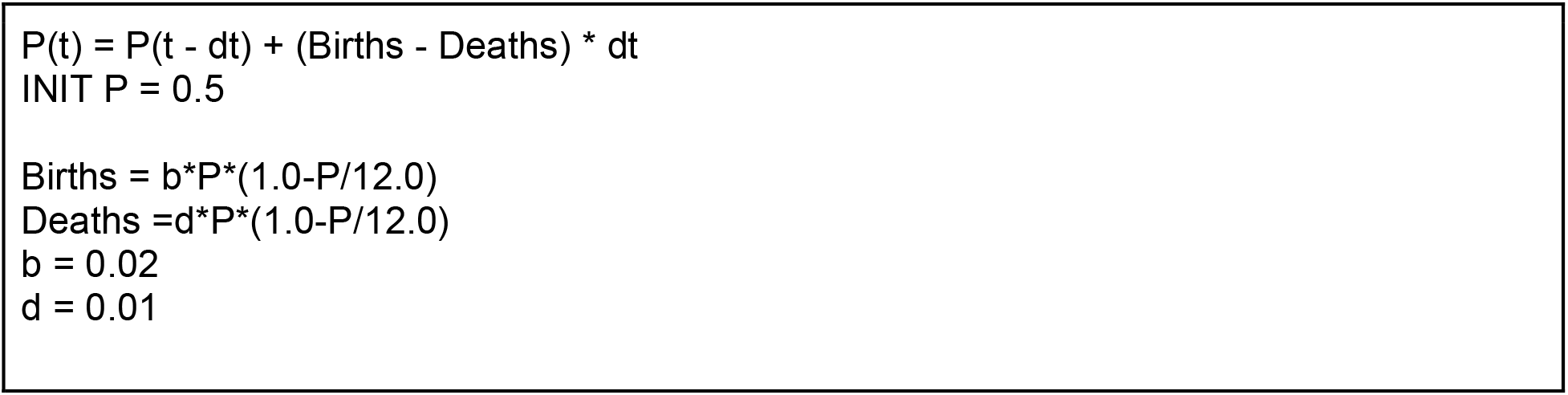
Stella equation of the graphical representation shown in Figure 1, which corresponds to the differential equation shown in Equation 1, and the values assigned to the parameters *b* and *d*.

**Figure 3.**
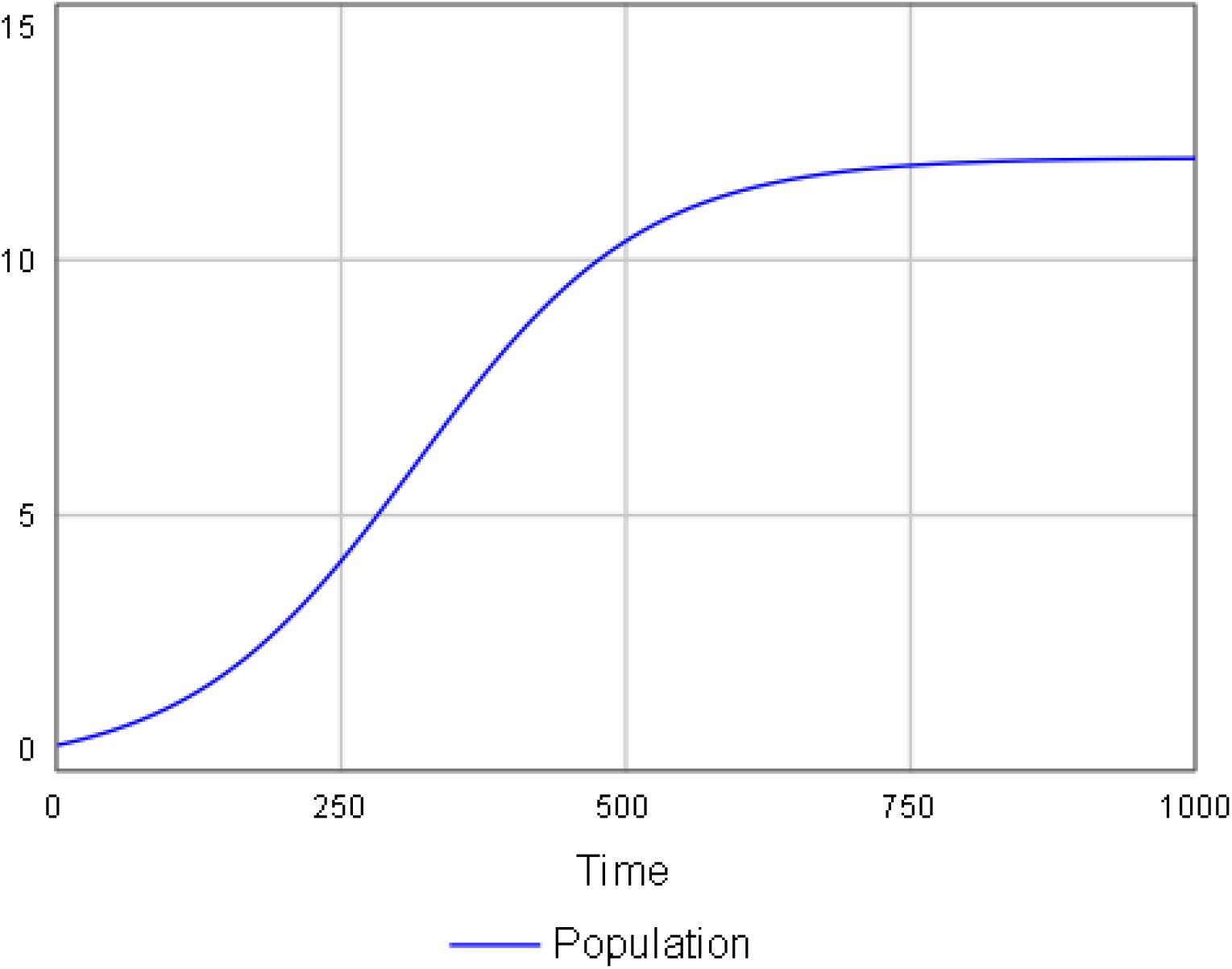
Solution in Stella of the model shown in Figures 1 and 2.

## 3 SRCompiler

SRCompiler analyses the equations of the model in Stella once they have been exported to text. In the first phase, a lexical analysis using FLEX (Fast Lexical Analyzer Generator) is applied to identify tokens of the Stella language. FLEX is a computer program for generating lexical analyzers written by Vern Paxson in C around 1987. Tokens take the form of (i) logical-mathematical characters as “=” or “+”, for instance; and (ii) the name of Stella functions, called Builtins, whose first letter is in capital. For example, the first line in Figure 2 corresponds to an ensemble of tokens defining Equation 1. The first token is the name of the function “P”; the second one is the symbol “(“; the third one is “t”, and so on. The second ensemble of tokens corresponds to the second line in Figure 2, where the initial value of the stock *P*(t) is declared. This ensemble of tokens is easily identifiable since it always starts with “INIT” followed by the name of a variable, which in turn is followed by an equal symbol “=” and a numerical value. The following sentences are different ensembles of tokens. The format in which the tokens are structured defines a formal language that allows the program to identify all the variables, equations, and functions. In this phase, a symbol table could be introduced to facilitate and speed up the tracking of the types of all variables and their memory assignment. However, we leave the implementation of such a table as a future improvement of the SRCompiler.

In the second phase, a syntax analysis is applied. To do so, we build a syntactical analyzer with a set of rules implemented in Bison, which is a specific software for that purpose. This set of rules is the grammar core used to recognize the valid input text files to be translated. Here, the syntax analyzer will identify any error regarding the order of the tokens. For example, Figure 4 shows the set of rules in *context-free grammar notation* for the example shown in Figure 2. To make this grammar understandable for readers not used to this notation, we have included a description of them in Table 1. The first two rules have the following structure:

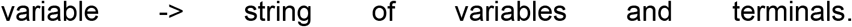

This notation defines a *variable* as a finite set of other symbols, each of which represents a language and *terminals*, which are symbols of the alphabet of the language being defined. The second rule, the one starting with “A ->”, classifies the token or set of tokens as one of the four pre-establised structures separated by the “|” symbol, which acts as a logical “or”. Such structures are as follows (Table 1):

1) The set of tokens starts with the name of a variable (“id”), followed by “(t)” (expressed as “pari tkT pard”), and then followed by “=” (“assig”), then the right part of the equation which starts with the same name of the variable (“id”), followed by “(t - dt)” (expressed as “pari tkT opas tkDT pard”), then the arithmetic operator (“opas”) which expresses the sum (“+”) or difference (“-”) and the last part where we can find a possible operation with variables or constants inside of parenthesis (“pari Op pard”) followed by another arithmetic operator.
2) The set of tokens starts with “INIT” (“tklnit”) followed by the name of the variable, then by “=” (“assig”), and then by a possible operation with a set of variables, constants or other outputs from Stella builtins (“F A”).
3) The set of tokens starts with the name of a variable (“id”), followed by “=” (“assig”), and then followed by a possible set of variables, constants or other outputs from stella builtins (“Op A”).
4) The variable “A” is empty.

**Table 1.**
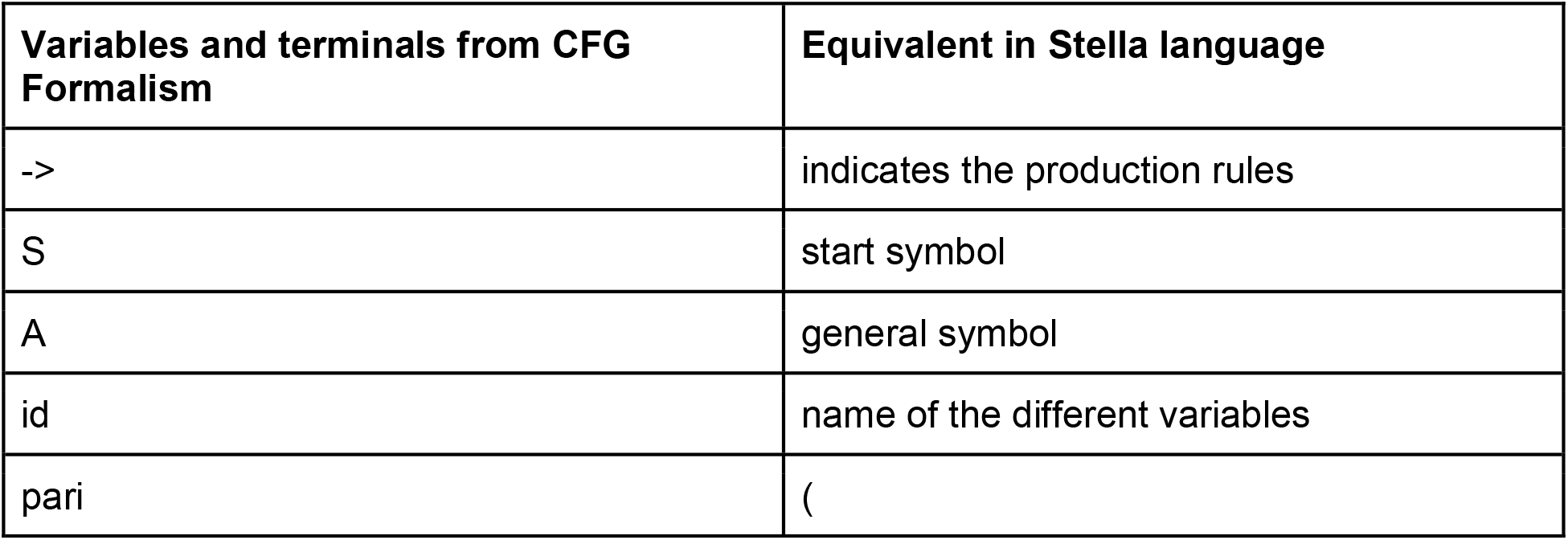
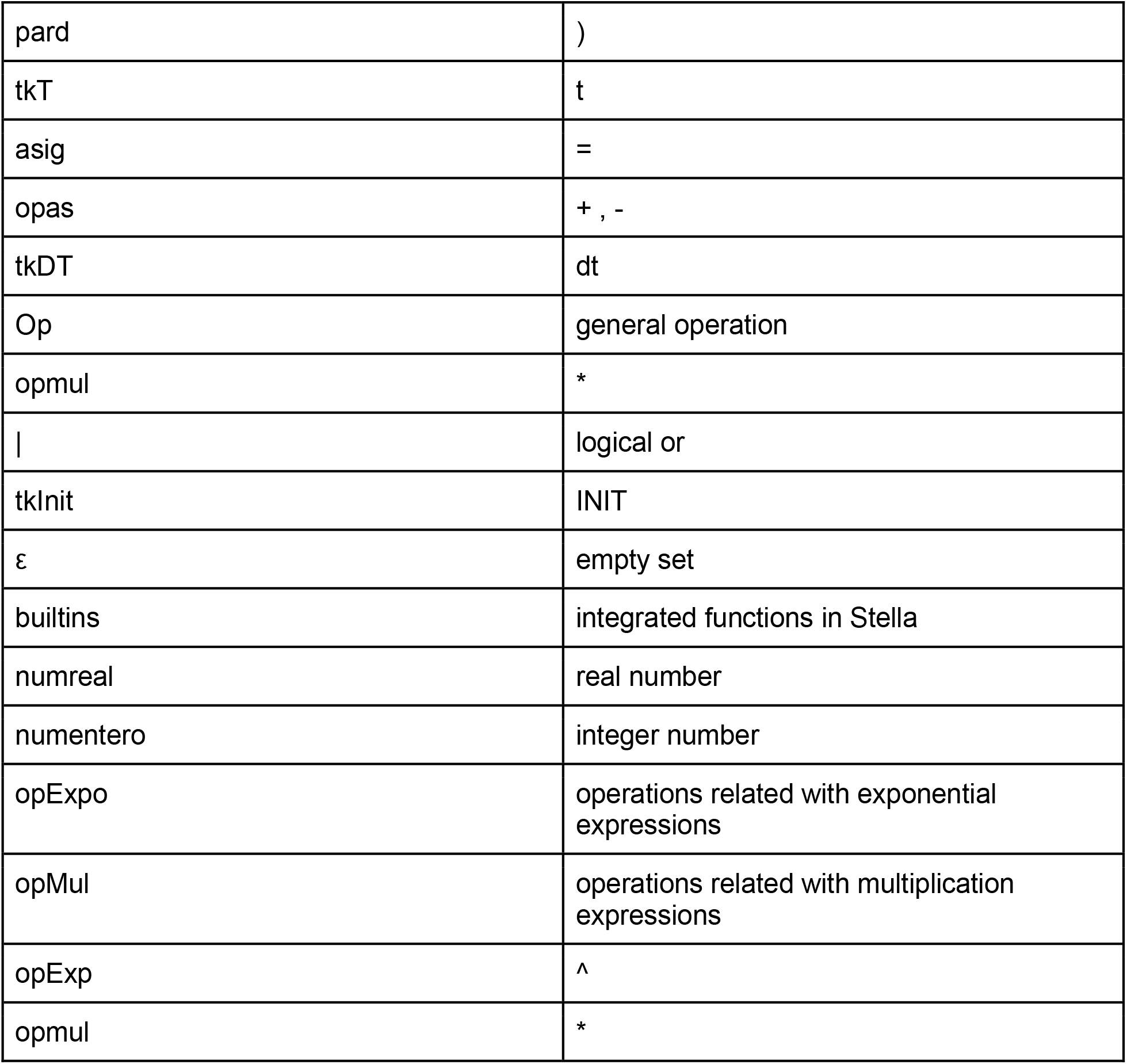
In the first phase the compiler uses the *lexical analyzer* which will divide the program into some meaningful strings which are known as a token (words at the left of the table), those tokens will be sended to the *syntax analyzer*.

**Figure 4.**
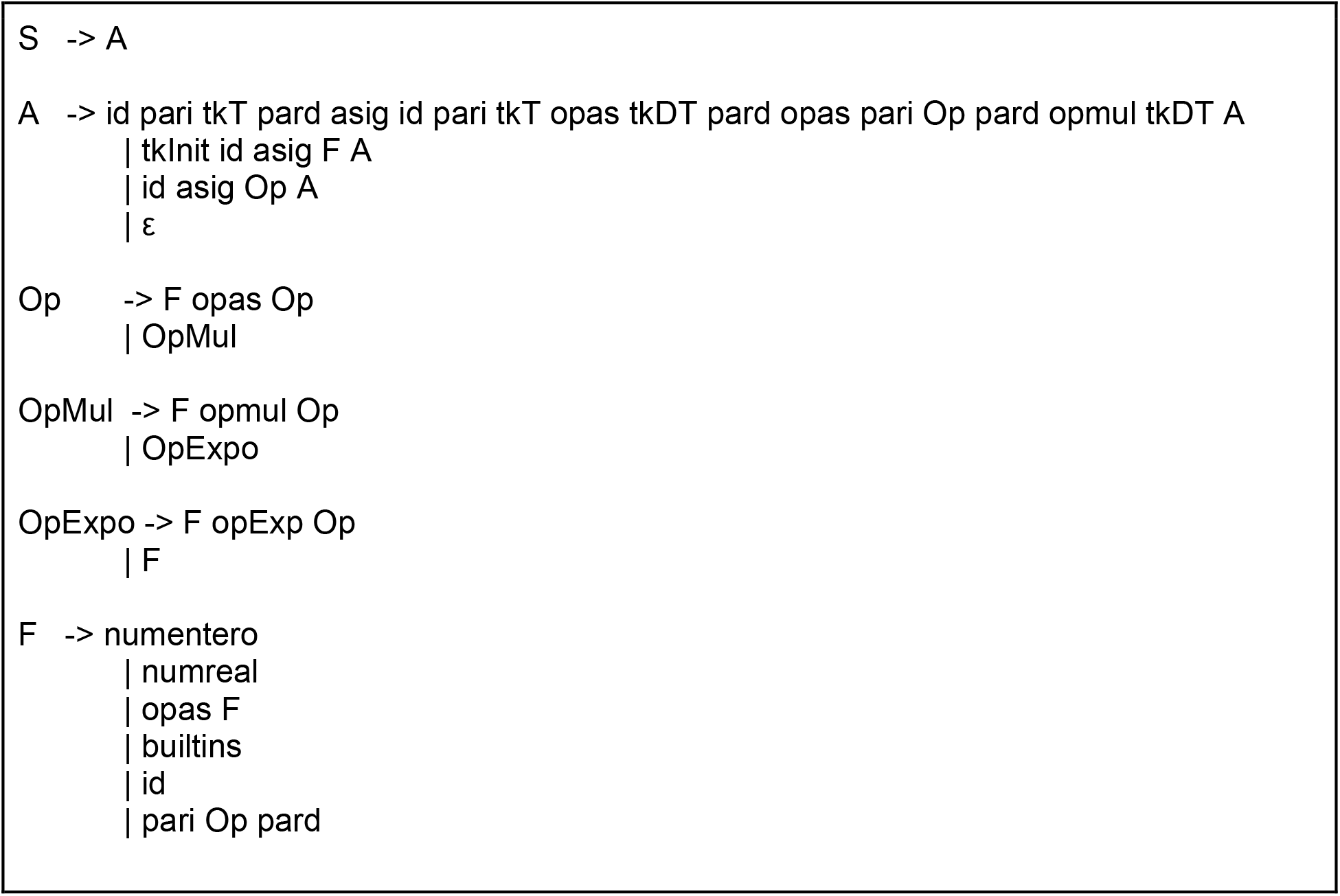
Grammar defined in the syntactical analyzer using *context-free grammar (CFG)*, which is a notation for describing languages.

The third rule, the one starting with “Op ->”, can be setted followed by an operation with variables, constants or outputs from builtins (“F”) with an arithmetic operator (“opas”), and with another (“Op”) rule. Or simply can be expressed by the fourth rule (“Opmul”).

The fourth rule, the one starting with “OpMul ->”, can be followed by an operation with variables, constants or outputs from builtins (“F”) with an arithmetic operator (“opmul”) which expresses the multiplication, and with another (“Op”) rule. Or simply can be expressed by the fifth rule (“OpExpo”).

The fifth rule, the one starting with “OpExpo ->”, can be followed by an operation with variables, constants or outputs from builtins (“F”) with an arithmetic operator (“opExp”) which expresses the exponentiation operation, and with another (“Op”) rule. Or simply can be expressed by any operation (“F”).

The sixth rule, the one starting with “F ->”, can be an integer number (“numentero”), a real number (“numreal”), an arithmetic operation (“opas”) followed by this same rule, an output from Stella’s builtins, the name of a variable (“id”), and finally an operation locked inside of parenthesis (“pari Op pard”).

In the third phase, a semantic analysis is applied. Here, the meaning of each syntactic construction and the semantic rules that must be obeyed are defined. For example, in our Stella code (Figure 2) the differential equation was written as:

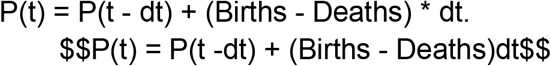

This variable declaration is always the same inside of the formalism established by Stella’s language. If such formalism is not respected as, for example, writing a similar expression with no parenthesis,

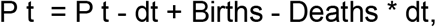

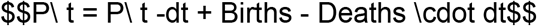

then the compiler will warn that a semantic error exists and it must be corrected according to the setted rules. Although no semantic errors are expected in the code from Stella Architect v. 2.1.4, they may appear in future versions due to the incorporation of new features, or in codes modified by the users by any reason.

Finally, if all the analyses are conducted without errors a R script is generated with the model translated from Stella. The variable names and their initial values in Stella are kept in the R script. In the first line of the R script the *deSolve* package is loaded to solve the dynamic model via the ODE function. Additionally, if any numerical values have been graphically defined by drawing in Stella, a CSV file is generated and then such values can be used in the R script.

Figure 5 shows the translation of the model in Figure 2 from Stella to R. The model has been encapsulated as a R function and the values of the converters and initial values are stored in the vector *parms* and *Y*, respectively.

**Figure 5.**
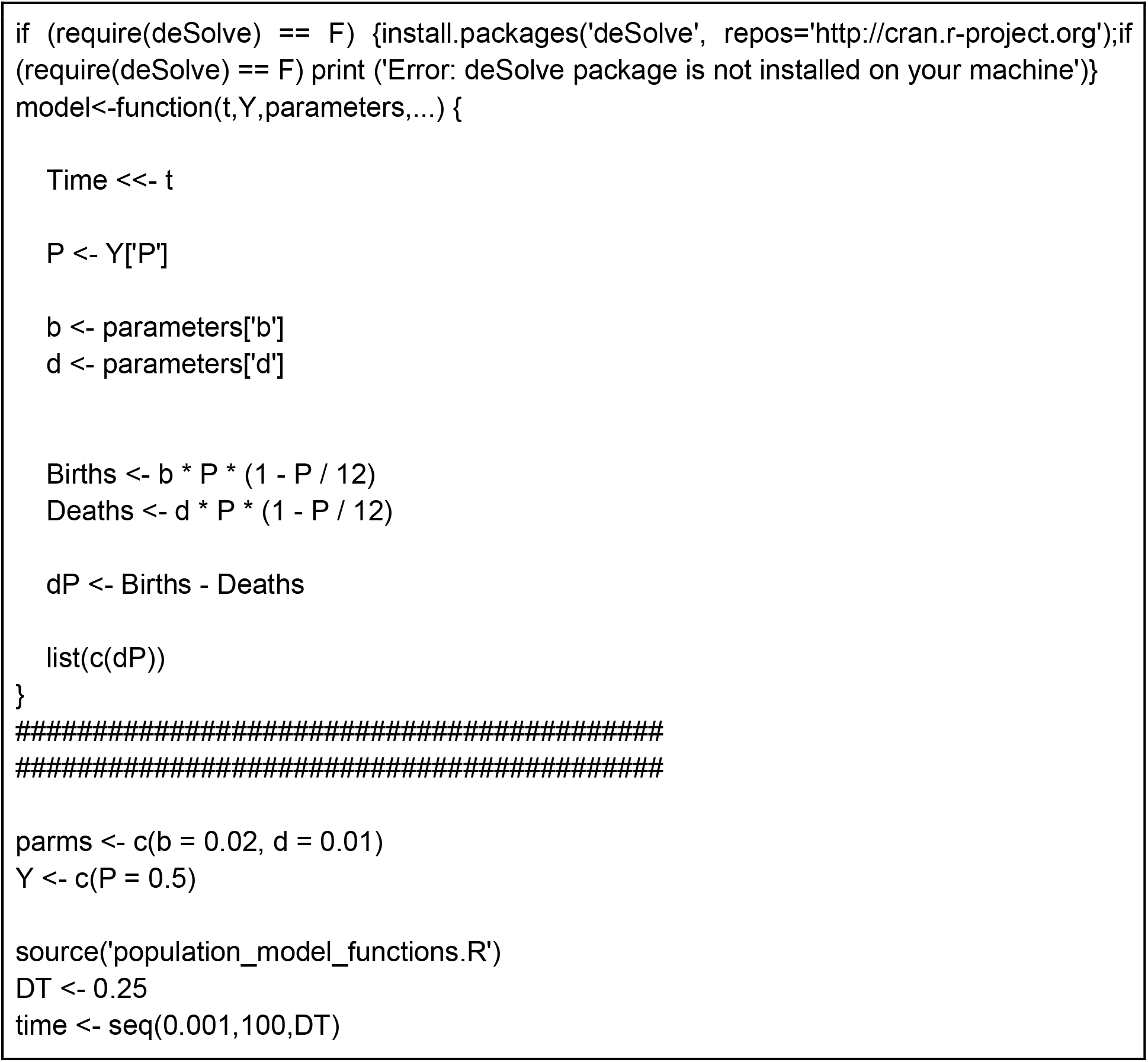
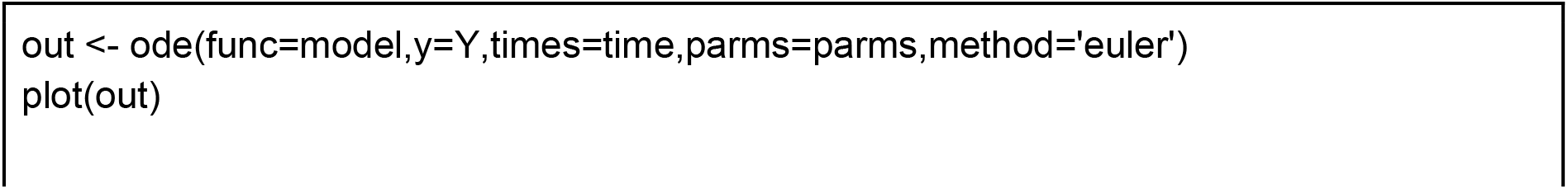
R script translated from the Stella model shown in Figure 2.

This first version of the SRCompiler is not able to translate all the Stella’s functionalities, but it is able to translate density-dependent dynamic models based on converters, flows and reservoirs, which are the fundamental elements of Stella. Besides, arithmetic operators (see Table 1) and some basic mathematical and trigonometric functions are also translated. Future versions of the SRCompiler will expand the supported Stella functions and mathematical operations.

## 4. Parameters optimization example

In this section, we will use an example to show the usefulness of the SRCompiler in, for example, population ecology. In particular, we will model a classical prey-predator dynamical system in Stella, whose parameters will be optimised to fit the snowshoe hare and the Canadian lynx populations in boreal forests of North America (Krebs, 2014). In abundance of hares, lynx mainly feed on them, which indicates a strong interconnection between their populations.

Let *H*(*t*) and *L*(*t*) represent the hare and lynx population at time *t*, respectively. These populations increase with births and decrease with deaths. Then,

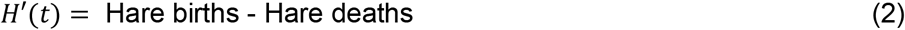

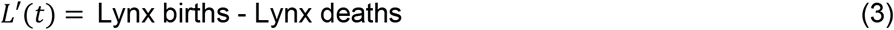

With unlimited resources, hare populations could grow indefinitely and hare births could be modelled as

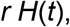

where *r* is the hare reproduction rate. However, resources are always limited and births should show a logistic growth limited by *k*, the hare carrying capacity of the ecosystem. Then, the hares births should be modelled as

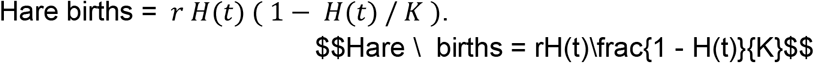

For the sake of simplicity, it is assumed that all hare deaths are produced by lynx predation, and lynx only consume hares which they have hunted. According to the first assumption, hare deaths can be modelled as follows (Tyson et al., 2010),

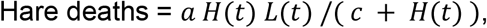

where *a* is the maximum lynx/hare predation rate, and *c* is the half-saturation constant, that is the hare density at which predation is at half *a*. The second assumption allows to model lynx births as

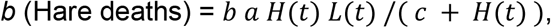

where *b* is the lynx/hare conversion rate. Finally, if *d* represents the lynx death rate, then Lynx deaths = *dL*(*t)*.

Then, equations 2 and 3 can be written as

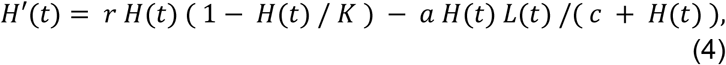

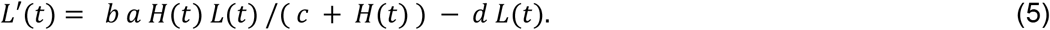

This model is a simplification of the real situation since hares are predated on by a number of predators such as golden eagles, great horned owls, coyotes or wolves, and lynx also predate on small rodents or red squirrels, as described by Stenseth et al. (1997). However, it is a classic example suitable to show the usefulness of the SRCompiler.

The model described in equations 4 and 5 can be written in Stella using a block diagram as in Section 2, as shown in Figure 6. The Stella representation of the model in text can be seen in Figure 7. The model is defined by 6 parameters: *a, b, c, d, k*, and *r*. Different values of the parameter will represent different prey-predator systems. In our case, we want to fit our model to the historical records from the Hudson’s Bay Company. Original data was published by MacLulich (1937), but we will use a digitised version from D. R. Hundley at Whitman College (http://people.whitman.edu/~hundledr/courses/M250F03/M250.html). During the process, the y-axis for lynx data was lost, producing a wrong amplification of them. Here, the lynx population has been rescaled to the original values by forcing it to reach a maximum of 6000 lynx in 1886 (Odum and Barrett, 2005). The annual population of hares and lynx have been estimated from the number of fur traded by trappers from 1844 to 1904, while data from 1905 to 1935 are derived from questionnaires (Figure 8; further details in Stenseth et al., 1997). It should be kept in mind that the reliability of these time series is limited by unknown external factors. For example, the number of caught animals may depend on the number of trappers, which, in turn, may depend on the purchase price of furs from one year to another.

**Figure 6.**
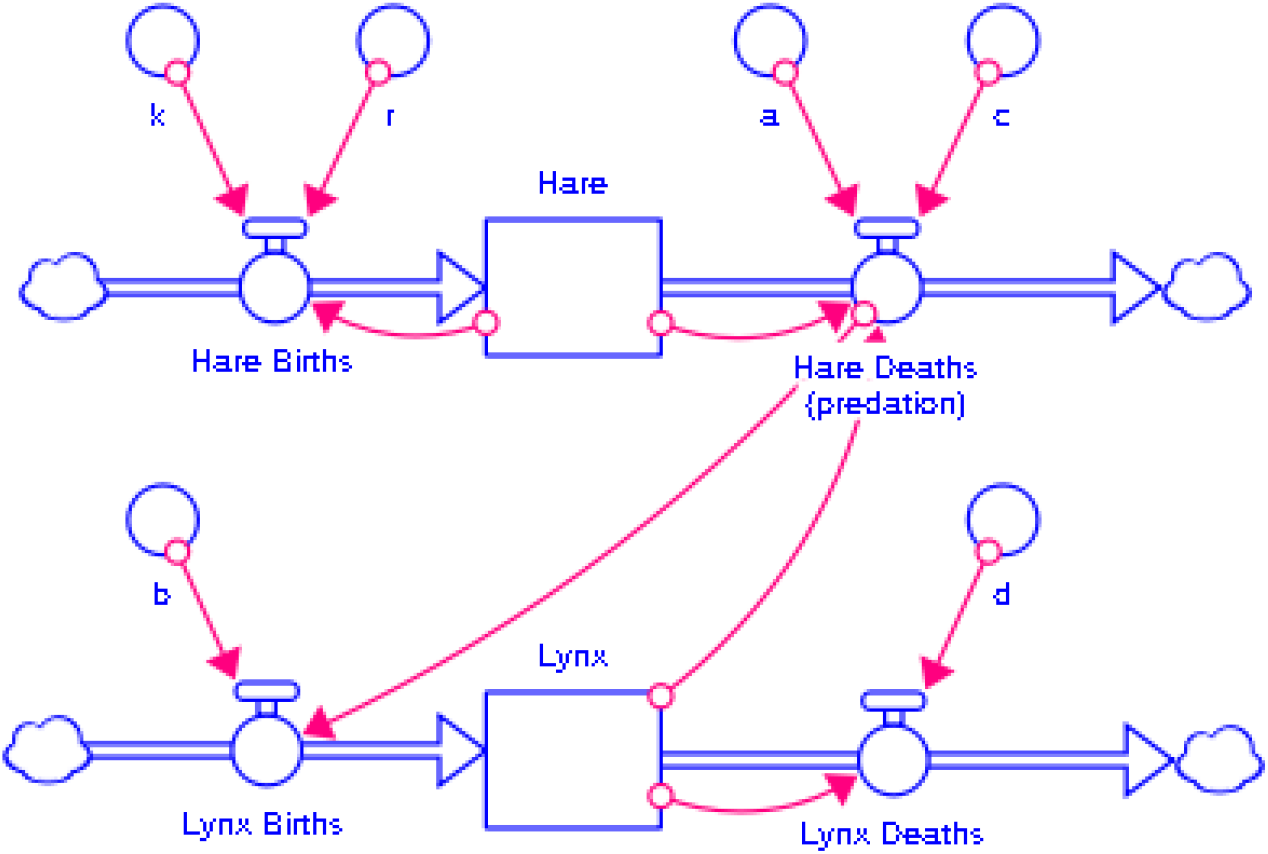
Hare-Lynx Model Population in Stella.

**Figure 7.**
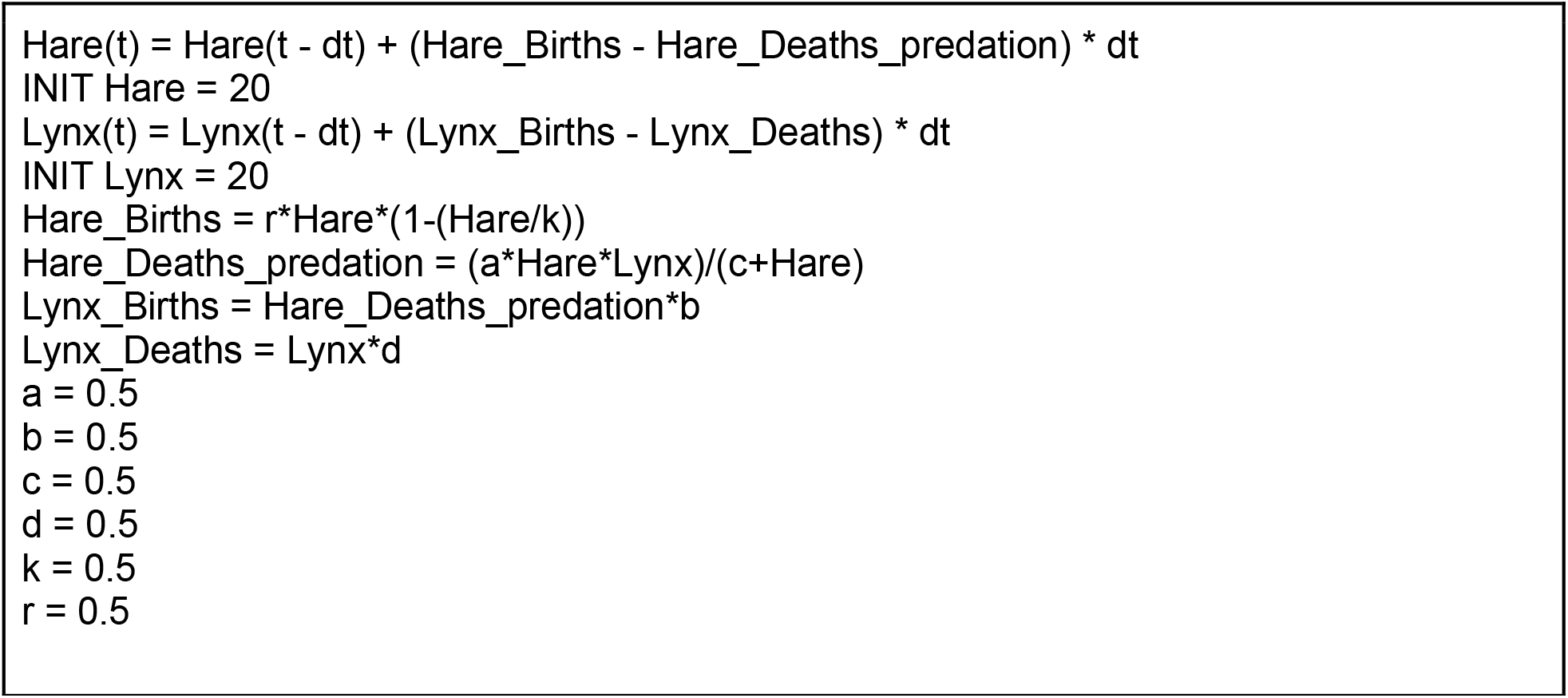
Stella equation of the graphical representation shown in Figure 6, which corresponds to the differential equation 4 and 5. The values of the parameters are equal to 0.5 because we need to declare them for the subsequent translation, although those values will be different after the optimization process.

**Figure 8.**
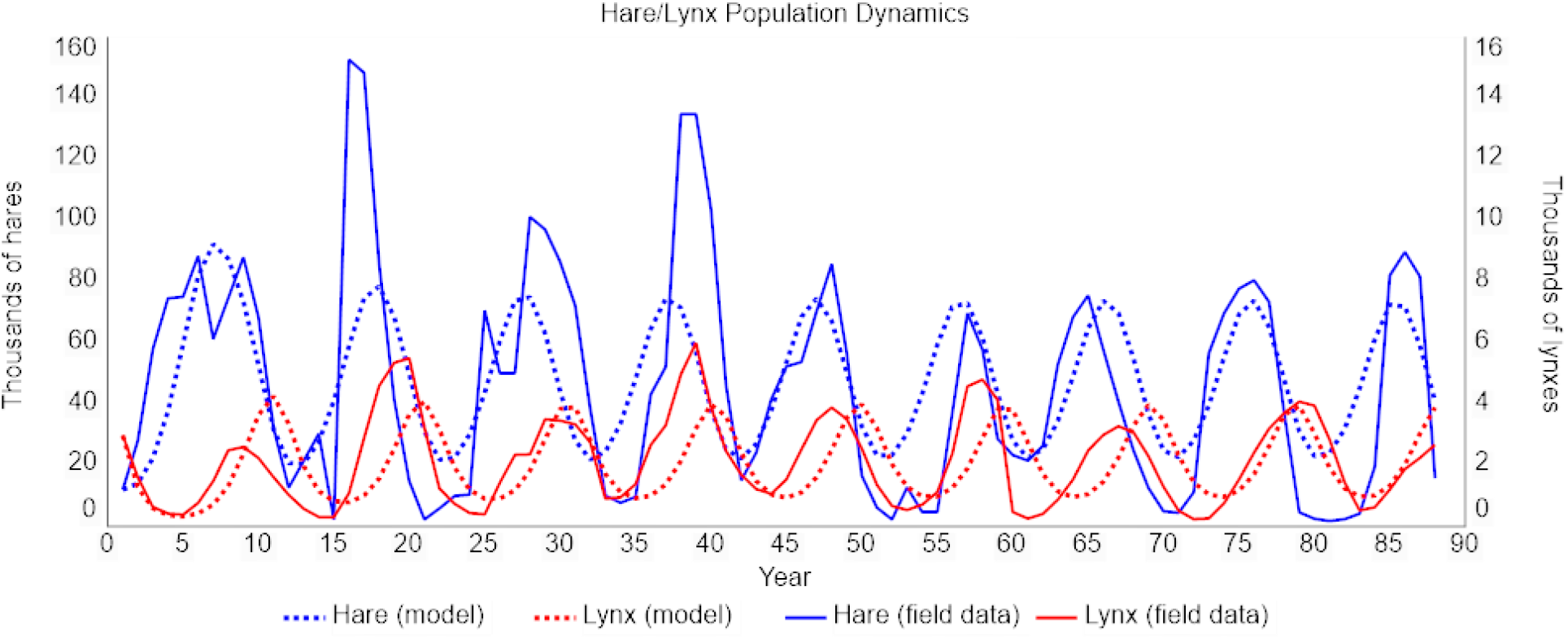
Historical (solid lines) and simulated (dashed lines) population of hares and lynxes. The simulation is done in Stella, based on equations 4 and 5 with the parameters optimised in R and shown in Table 2. The y axis is not the number of individuals but a proxy proportional to the actual population.

The problem now is to figure out the value of the parameters to get a good approximation between the lynx/hares population simulated by the model and the historical data. To do so, we translated the Stella model to R using the SRCompiler. Then, we used the *SCEoptim* function from the *SoilHyP* package to optimise the parameters. This function is a global optimization algorithm based on The Shuffled Complex Evolution (SCE-UA) method developed at the University of Arizona (Naeini et al., 2019). The fixed rank for the parameters was

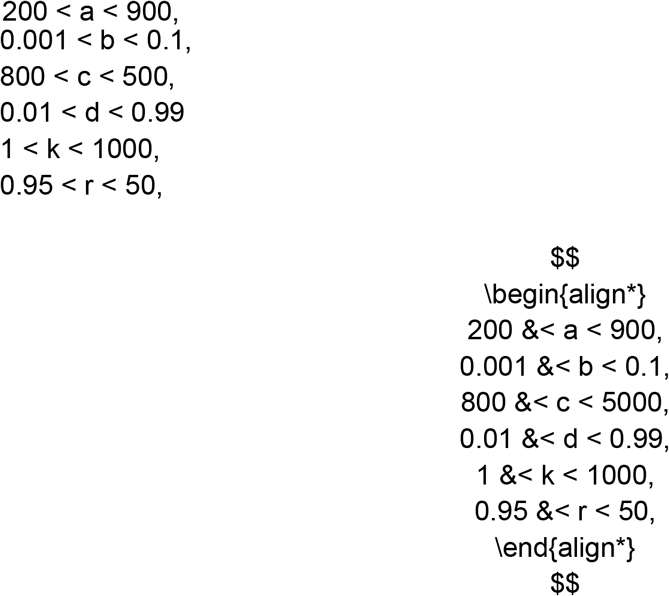

and the options of the algorithm were 30 for number of complexes and 3 for elitism. The obtained parameters are shown in Table 2. Once they are introduced in Stella, the model output agrees quite well with the historical data (Figure 8). The number of cycles is well reproduced in both lynx and hare populations, as well as the delay between both populations. The timing of the maxima and minima is quite reasonable too. As it was expected, the model reproduces a steady situation, but not the amplitude variability of the cycles.

**Table 2.** Parameters optimised in R to fit the model (Equations 4 and 5) simulation to historical data of hare and lynx. Optimization was implemented using the Shuffled Complex Evolution (SCE-UA) (Naeini et al., 2019). The model in R is a translation of the Stella model (Figures 6 and 7) by SRCompiler.

| Parameter | Symbol | Value |
| --- | --- | --- |
| Maximum lynx/hare predation rate | $a$ | 900 |
| Lynx/hare conversion | $b$ | 0.714 |
| Half-saturation constant | $c$ | 3492 |
| Linx death rate | $d$ | 0.791 |
| Hare carrying capacity | $k$ | 105 |
| Hare reproduction rate | $r$ | 1.06 |

## 4. Discussion

In this study we have developed a translator from Stella to R software, named SRCompiler, which has been used to estimate the parameters of a classic predator-prey to fit the historical time series of the snow hare and lynx in Canada from the Hudson Bay Company. The translator is based on compiler design and defines a formal language, which makes it more versatile, scalable and expandable than a previous translator based on a text procedure (Naimi and Voinov, 2012). This is so because the latter is less robust to changes when the source code from Stella gets an update changing names of words or add additional information related with the dynamic model. SRCompiler is freely available and can be downloaded from github (https://github.com/hmpp91/SRCompiler).

As well as other softwares such as Vensim or Powersim, Stella provides a powerful and friendly environment to design dynamic models, which can be used by people that are not comfortable with the mathematical expressions of differential equations, or by mathematical and population ecology experts that appreciate the advantages of a visual design of complex dynamic models. In any case, once the model has been built, it is not easy, and in some cases not possible, to analytically optimise the parameters of the model in Stella to better fit the simulation to real data. However, this is a crucial point to make a theoretical model become a realistic one, or at least as realistic as possible. Once the model reproduces the experimental data, it is ready to perform experiments that help us understand the dynamic system. For example, the evolution of the system could be studied from variations in the parameters, which represent different possible scenarios, shedding light on the possible situations that will keep the system in balance or will collapse it. This “What if” analysis is important even if the populations are only proportional to the real populations as in the hare-lynx example.

The importance of the SRCompiler lies precisely in the determination of model parameters since it allows translating the Stella model to R software, where several parameter optimization tools are freely available. The process would be as follows: (1) the Stella model is translated to R; (2) the parameters are optimised in R within a range of biologically meaningful values to fit the experimental data; (3) the values of the calculated parameters are assigned to each variable in the Stella model. Then, the Stella model becomes more realistic as the simulated values are as close as possible to the data.

The optimization can still be improved. In the classic hare-lynx population example we have obtained a set of 6 constant parameters, reaching a reasonably good approximation to the data. However in population ecology or other biological disciplines, variability over time is a characteristic of each variable, which implies time-variable coefficients. The SRCompiler is also useful in this situation, as we plan to show in a further study together with the “what if” scenario analysis. The latter could be relevant to analyse the influence in the system of increased hunting intensity, decreased food for hares, a one-off event such as an epidemic, etc.

This methodology of optimising population parameters would allow to revisit population models and to recalculate solutions, therefore increasing the ability to compare different scenarios (e.g. What if scenarios). We believe that the science of population biology, applied to both conservation and sustainable exploitation (e.g. predator-prey or competing species, fisheries, epidemics dynamics, etc) has much to benefit from the use of these new tools, which enables the application of a wide range of mathematical analysis to Stella users.

## Acknowledgements

We are grateful to the Fundació Baleària (https://fundaciobalearia.org/), Marina El Portet de Dénia (https://marinaelportet.es/), Marina de Dénia (https://marinadedenia.com/) and Varadero Port Dénia (https://www.portdenia.com/) for the support of the MarLabUA-Dénia.

